# Neuropeptidergic regulation of neuromuscular signaling in larval zebrafish alters swimming behavior and synaptic transmission

**DOI:** 10.1101/2024.01.12.575339

**Authors:** Holger Dill, Jana F. Liewald, Michelle Becker, Marius Seidenthal, Alexander Gottschalk

**Affiliations:** Buchmann Institute, Goethe University, Max-von-Laue-Strasse 15, D-60438 Frankfurt, Germany; Institute for Biophysical Chemistry, Goethe University, Max-von-Laue-Strasse 9, D-60438 Frankfurt, Germany

**Keywords:** Synaptic transmission, neuromodulation, neuropeptides, zebrafish, homeostatic scaling, tachykinin, electrophysiology, behavioral analysis

## Abstract

The regulation of synaptic transmission is crucial for plasticity, homeostasis and learning. Chemical synaptic transmission is thus modulated to accommodate different activity levels, which also enables homeostatic scaling in pre- and postsynaptic compartments. In nematodes, cAMP signaling enhances cholinergic neuron output, and these neurons use neuropeptide signaling to modulate synaptic vesicle content. To explore if this mechanism is conserved in vertebrates, we studied the involvement of neuropeptides in cholinergic transmission at the neuromuscular junction of larval zebrafish. Optogenetic stimulation by photoactivated adenylyl cyclase (bPAC) resulted in elevated locomotion as measured in behavioural assays. Furthermore, post-synaptic patch-clamp recordings revealed that in bPAC transgenics, the frequency of miniature excitatory postsynaptic currents (mEPSCs) was increased after photostimulation. These results suggested that cAMP-mediated activation of ZF motor neurons leads to increased fusion of SVs, consequently resulting in enhanced neuromuscular activity. We generated mutants lacking the neuropeptide processing enzyme carboxypeptidase E (cpe), and the most abundant neuropeptide precursor in motor neurons, tachykinin (tac1). Both mutants showed exaggerated locomotion after photostimulation. *cpe* mutants exhibit lower mEPSC frequency during photostimulation and less large-amplitude mEPSCs. In *tac1* mutants mEPSC frequency was not affected but amplitudes were significantly smaller. Exaggerated locomotion in the mutants thus reflected upscaling of postsynaptic excitability. *cpe* and *tac1* mutant muscles expressed more nicotinic acetylcholine receptors (nAChR) on their surface. Thus, neuropeptide signaling regulates synaptic transmitter output in zebrafish motor neurons, and muscle cells homeostatically regulate nAChR surface expression, compensating reduced presynaptic input. This mechanism may be widely conserved in the animal kingdom.

## INTRODUCTION

Proper function of the animal nervous system depends on secretion of chemical neurotransmitters such as acetylcholine (ACh) from the presynaptic terminal into the synaptic cleft, and binding of these transmitters to their respective receptors in the postsynaptic compartment. The molecular mechanisms behind this dynamic process and how it is regulated are still not fully understood [1]; however, synaptic efficacy is often controlled by neuromodulators [2, 3]. Neurotransmitters are stored in small organelles termed synaptic vesicles (SVs) from which they are released by the neuron through exocytosis. Fusion of SVs with the plasma membrane at the active zone of the synapse is triggered by depolarization and the resulting increase in Ca^2+^ concentration. SV exocytosis is promoted in response to intracellular signaling via cyclic adenosine monophosphate (cAMP) and protein kinase A (PKA) signaling, mediated by diverse synaptic PKA targets that induce SV mobilization and SV priming, such that increases in Ca^2+^ concentration have a higher chance of triggering SV fusion events [4]. Some known PKA targets are synapsin [5], tomosyn [6], Rim1 [7], ryanodine receptor (RyR) [8], cysteine string protein [9], snapin [10], complexin [11], and SNAP-25 [12]. In general, it is beneficial if synaptic transmission can be modulated, to adapt to the demands of the current situation, or to alter synaptic efficacy in the long term. On the presynaptic side, modulation of transmission can be mediated by altering the response of the presynaptic machinery to changes in the Ca^2+^ concentration, thus altering the rate at which SVs are mobilized from the reserve pool [13] or varying the (quantal) content of neurotransmitter per SV [14]. Furthermore, neuromodulators can affect presynaptic transmission, e.g. by altering the number of SVs being produced [3] . Postsynaptic modes of regulating signal efficacy have also been described, for example changes in the amount of postsynaptic transmitter receptors and their interaction with scaffolding proteins, membrane resistance or other modes of excitability, like upregulation of voltage-gated ion channels [15–17].

Although synaptic architecture varies between cell types and species, the ultrastructure of synapses, the composition of SVs and the ‘pools’ into which they are organized are conserved between invertebrates and vertebrates including zebrafish (*Danio rerio*) [18]. Structures at neuromuscular junctions (NMJs), the interface between cholinergic motor neurons and skeletal muscle fibers, have been observed in zebrafish larvae using ultrastructural imaging techniques. As in other organisms, they comprise SVs, dense core vesicles (DCVs) containing neuropeptides, as well as clathrin-coated vesicles [19]. Like other peptide hormones, neuropeptides, which are stored in and released from DCVs, originate from longer precursor proteins which are packaged into vesicles in the *trans*-Golgi network, which are subsequently acidified. Neuropeptide precursor proteins are converted into small bioactive peptides by pro-protein convertase cleavage and C-terminal trimming by carboxypeptidase E (CPE) [20]. DCVs are then transported to the synaptic terminals for release [21, 22]. In zebrafish, remarkably diverse functions have been associated with neuropeptides. For example, the expression levels of parathyroid hormone 2 (*pth2*) were shown to mirror the presence of conspecifics and also the density of the swarm [23]. Furthermore, galanin (*galn*) expression in the brain is essential for the generation of color patterns by self-organization of pigment cells [24], while tachykinins (*tac*) are broadly expressed in the central nervous system and have a supposed role as neuromodulators [25, 26].

The development of optogenetic tools has revolutionized the field of neuroscience by enabling researchers to manipulate specific neurons or neuronal populations. Today, optogenetic methods have become standard in many model organisms and thus, optogenetic tools to influence neuronal activity have also been established in zebrafish [27, 28]. Microbial rhodopsins for de- and hyperpolarization were used to study the function of sensory neurons [29] or central pattern generators [30] as early as day one of embryonic development, allowing to achieve calibrated light-induced currents and behavior in different sets of neurons, among them motor neurons [27]. In addition to light-activatable ion channels and pumps, other optogenetic tools have emerged in recent years. One class of tools for optogenetic manipulation are photoactivated adenylyl cyclases (PACs) such as bPAC, mediating light-dependent synthesis of the second messenger cAMP for precise temporal and spatial control and acute tuning of intracellular cAMP concentrations [31–34]. bPAC was shown to be enzymatically active in early zebrafish embryos [35] and has been applied to analyze the role of cAMP signaling in axonal regeneration [36] or to induce swimming behavior by activating hindbrain reticulospinal V2a neurons [37].

We have previously analyzed the effects of cAMP generation in presynaptic terminals of cholinergic motor neurons in the nematode *Caenorhabditis elegans*. This uncovered an as yet unknown role of neuropeptides in the regulation of synaptic transmission and plasticity [14], which, apart from increasing the rate of SV release, also affected the filling state of SVs with ACh. Thus, synaptic transmission can be controlled by two regimes: Depolarization and Ca^2+^ concentration control the acute fusion of SVs, while cAMP and peptidergic signaling alter the transmitter content per SV. This way, in addition to network activity of central pattern generators, motor neurons can integrate pathways of neuromodulation in different systemic states.

Expression of bPAC and the consequences of increased synaptic cAMP levels have not been described in zebrafish motor neurons to our best knowledge. In this study we explored whether the dual control of neuronal transmission, as described in *C. elegans* [14], is conserved also in vertebrates, specifically in zebrafish. We expressed bPAC in spinal motor neurons and photostimulated larvae at different developmental stages. This evoked exaggerated, but coordinated locomotion, as well as increased SV fusion rates and miniature excitatory postsynaptic currents (mEPSCs). We generated deletion alleles of the main neuropeptide processing enzyme, *cpe*, as well as of the most abundantly expressed neuropeptide in cholinergic motor neurons, *tac1*. Both mutants showed enhanced behavioral effects in response to bPAC stimulation, which likely occurred through postsynaptic compensation of presynaptic defects, like smaller mEPSC amplitude and lower SV release rates. This postsynaptic compensation involves increased expression of nicotinic acetylcholine receptors (nAChRs) in the neuromuscular endplates. Thus, mechanisms of cholinergic neuromodulation by neuropeptides appear to exist also in zebrafish, even though the details seem to differ between vertebrates and invertebrates.

## MATERIALS AND METHODS

### Animal husbandry and housing

Adult zebrafish (*Danio rerio*) were maintained in groups inside 6 liter tanks (5-7 fish per liter) located in a circulating water system (Zebcare, Nederweert, The Netherlands) with a 14 h / 10 h light / dark cycle and in accordance with FELASA guidelines [38]. Prior to all optogenetic experiments zebrafish embryos were kept in E3 medium in an incubator at 28°C and total darkness. Developmental stages of embryos were determined as described [39]. All experiments employing animals were conducted according to the European Directive 2010/63/EU on the protection of animals used for scientific purposes and the animal research board of the State of Hessen (animal protocol approval number V54-19c20/15-FR/1017, V54-19c20/15-FU/Anz. 1018, V54-19c20/15-FR/1019, and V54-19c20/15-FR/2003).

### Molecular biology

To obtain the Tol2-mnx1En:bPAC-egfp-polyA plasmid (pHD5) the coding sequence of bPAC was amplified with primers bPAC-attB1-F (ggggacaagtttgtacaaaaaagcaggctgcgccaccatgatgaagcggctggtgta) and bPAC-attB2-R (ggggaccactttgtacaagaaagctgggtagtacgtccgcggcttgtcgttt) and subjected to a Gateway™ BP-reaction (Thermo Fisher Scientific, Waltham, USA) with donor vector pDONR221 resulting in the middle entry clone pME-bPAC (pHD1). The middle entry clone (pHD1) was used in an Gateway™ LR-reaction together with the p5E-mnx1En [40], p3E-polyA and the pDestTol2CG clones [41]. Furthermore, the plasmid Tol2-UAS-ChR2-P2A-dTomato-polyA, a gift from Dr. Soojin Ryu (Johannes Gutenberg University, Mainz) was used to generate the Tg[*UAS:ChR2-P2A-dTomato*] zebrafish line after sequence validation.

Templates for transcription of antisense *in-situ* probes were amplified from 24 hpf cDNA by using specific PCR primers cpe_F_2 (CCCATCTCAAACGCCTCTGT), cpe_R_2 (ATAAGTCTGGACGCAGTGCC), tac1_F (GATGGGGAAACGGTCCTCTG) and tac1_R (GCGCAGGACTGTCGGTATTA). PCR products were cloned into the pCRII-TOPO vector (Thermo Fisher Scientific, Waltham, USA). Resulting plasmids cpe_Ribo2 (pHD18) and tac1_Ribo (pHD25) were transcribed with Sp6 or T7 RNA polymerase in a digoxigenin labeling reaction.

CRISPR / Cas9 target sites were identified and sgRNAs designed as described in [42]. The following DNA oligos were used together with the DR274 plasmid to construct templates for sgRNA transcription: cpe_Oligo_1-1 (TAGGACAGCGCAGAAAACAGGA), cpe_Oligo_1-2 (AAACTCCTGTTTTCTGCGCTGT), cpe_Oligo_3-1 (TAGGGTCGCGAGCTGCTCGTGC), cpe_Oligo_3-2 (AAACGCACGAGCAGCTCGCGAC), tac1_Oligo_1-1 (TAGGAAGTAACTAAAGTTTAGA), tac1_Oligo_1-2 (AAACTCTAAACTTTAGTTACTT), tac1_Oligo_2-1 TAGGATTTATTTAACATGCTTA) and tac1_Oligo_2-2 (AAACTAAGCATGTTAAATAAAT).

### Whole mount *in-situ* hybridization

Whole mount *in-situ* hybridization was carried out as described previously [43]. In brief, after PFA fixation embryos were stored in MetOH and subsequently rehydrated. Embryos were treated with Proteinase K (5 μg/ml) according to the developmental stage. Proteinase K treatment was followed by refixation in 4% PFA for 20min. Unspecific RNA binding sites were blocked with Torula yeast RNA (Sigma-Aldrich, St. Louis, USA) before incubation with DIG-labelled RNA probes. Hybridization was carried out at 65°C overnight. To detect specifically bound RNA probes, embryos were incubated with AP-conjugated-anti-DIG antibodies (Roche, Basel, Switzerland) and stained with NBT/BCIP (Roche, Basel, Switzerland).

### Microinjection of zebrafish embryos

In order to generate transgenic zebrafish lines, plasmid DNA was diluted to a final concentration of 12.5 ng/µl in water and approximately 0.5 nl were co-injected with *in-vitro* transcribed *Tol2* mRNA (12.5 ng/µl) into 1-cell stage embryos. 2 dpf embryos were scored for expression of the *cmlc2:EGFP* transgenesis marker [41] and kept to establish the F0 generation.

To generate gene knockouts a pair of sgRNAs per gene (12.5 ng/µl each) were coinjected with Cas9 protein (Integrated DNA Technologies, Coralville, USA) as recommended by the manufacturer. After microinjection zebrafish embryos were kept in E3 medium in an incubator at 28°C. Primary injected F0 animals were mated with AB wildtype partners to obtain the F1 generation.

### Genotyping of adult zebrafish

Fin biopsies were taken from adult zebrafish and genotyping PCRs for the respective *cpe* or *tac1* knockout alleles performed on tissue lysates using the primers cpe_Geno_F (caaatatatgtgacccgttcgtc), cpe_Geno_R (ggcgatcctccattattgattgg), tac1_Geno_F3 (gctcacctcctctgacgtaa) and tac1_Geno_R2 (TGTGAAATGTCACTAACTTTGTTGC) with Phusion DNA Polymerase (Thermo Fisher Scientific, Waltham, USA).

### Genotyping of live zebrafish embryos

Genotyping of 48 hpf embryos was done as described in [44]. The chorion was removed manually and embryos loaded onto a microfluidic chip in E3 followed by agitation on a base unit (wFluidx, Salt Lake City, USA). 10 µl of E3, containing genetic material, was used as template for genotyping PCRs as described above. Subsequently, embryos were transferred to 24-well plates with fresh E3 to recover.

### Quantitative PCR

Total RNA was recovered from adult brain tissue by chloroform/phenol extraction using Trizol (Thermo Fisher Scientific, Waltham, USA) with subsequent isopropanol precipitation. cDNA was synthesized from 700 ng of total RNA with SuperScript™ II Reverse Transkriptase (Thermo Fisher Scientific, Waltham, USA) and oligo(dT) primers. Quantitative PCR was performed using the iTaq™ Universal SYBR® Green Supermix (Bio-Rad, Hercules, USA) in a CFX Connect Real-Time cycler (Bio-Rad, Hercules, USA) as described in the respective manuals. The average Ct values of replicates were normalized to gapdh to obtain ΔCt values. For relative expression analysis ΔΔCt values were calculated as described in [45]. The following primer pairs were used for quantitative PCR: cpe_qPCR_F2 (GGTCAACTACATAGAGCAGGTTCA), cpe_qPCR_R2 (CCAACAAGCGCCAGTAGTCA), tac1_qPCR_F2 (ATCGGTCTGATGGGGAAACG), tac1_qPCR_R2 (ACGACTCTGGCTCTTCTTGG), gapdh_qPCR_F (CAGGCATAATGGTTAAAGTTGGTA) and gapdh_qPCR_R (CATGTAATCAAGGTCAATGAATGG).

### α-Bungarotoxin Staining

Larvae (4 dpf) were fixed with 4% paraformaldehyde in PBST at 4 °C over night. Heads were cut off to allow diffusion into the tissue and larvae were briefly digested with collagenase (1 mg/ml) for 15 minutes. Staining of acetylcholine receptors was performed with 10 µg/ml CF®488A conjugated α-Bungarotoxin (Biotrend, Cologne, Germany) in PBST complemented with 10% FCS and 1% DMSO.

### Confocal laser scanning microscopy and image analysis

For imaging of chemically fixed samples, embryos or larvae were treated with 4% formaldehyde (Polysciences, Warrington, USA) in PBST at 4 °C over night and mounted in 1.5 % low melting point agarose (Thermo Fisher Scientific, Waltham, USA). *In-vivo* imaging was performed on larvae embedded in 1.5 % low melting point agarose in E3 medium supplemented with 4.2 g/l MS-222 (Sigma-Aldrich, St. Louis, USA). Confocal imaging was done with a Zeiss LSM 780 microscope using Plan-Apochromat 10x/0.3 Air or Plan-Apochromat 20x/0.8 Air objectives. Images were processed and particle analysis of α-Bungarotoxin stained samples was done with ImageJ [46].

### Zebrafish behavioral assays

For all behavioral experiments all embryos were kept in E3 medium and total darkness at 28°C previous to the assay. Animals were selected for the respective fluorescence marker with a Leica MZ16F stereomicroscope the day before the experiment. All assays were conducted at room temperature in pre-warmed E3.

### Evoked tail-coiling

At 24 hpf ∼15 – 20 embryos were transferred to a 6 cm petri dish and the tail-coiling behavior monitored for 30 seconds before and 30 seconds during the exposure to blue LED light (470 nm, 0.1 mW/mm^2^). In all behavioral assays, to stimulate bPAC as well as the wildtype control continuous blue light was applied. To activate ChR2 20 Hz light pulses (25 ms pulse length) were used. Videos were captured with an infrared background illumination at 30 frames per second for 1 minute on a similar video recording assembly as described in [47]. Spontaneous and evoked tail movements (STMs) were identified by a previously described MATLAB application [48].

### Tail-beat-assay with immobilized larvae

Zebrafish larvae at an age of 4 dpf were mounted in 1.5 % low melting point agarose (Thermo Fisher Scientific, Waltham, USA) and covered with E3. Agarose around the tail was removed with a scalpel and the tail remained free to perform tail beat movements. Animals were kept in the dark for 10 seconds followed by a 10 second blue light illumination period (460 nm LED, 0.1 mW/mm^2^). Videos were recorded with an Evolve Delta camera (Teledyne Photometrics, Tucson, USA) at a frame rate of 60 fps on a Zeiss Observer Z1 microscope equipped with an EC Plan-Neofluar 1x/0.025 M27 objective and red background illumination. The maximum angels between body axis and tail tip were extracted manually from minimum intensity projections in ImageJ.

### Swimming assay

For analysis of swimming behavior, single animals at 4 dpf were transferred to an agarose arena with a diameter of 3 cm filled with E3 medium and adapted for 2 minutes under dark conditions. Videos were taken as described above with a frame rate of 30 fps. Video files of swimming behavior were recorded for 20 seconds. After a dark phase of 5 seconds, continuous or 20 Hz blue light (470 nm, 0.1 mW/mm^2^) was applied from an LED ring for 10 seconds. Larvae were tracked with a custom macro in ImageJ (version 1.52). For each video frame pigmentation was smoothed by median filtering, animals separated from the background by thresholding and center of mass identified to obtain X- and Y-coordinates (as described in [49] and [50]). Distance travelled between consecutive video frames was calculated in pixels and converted into swimming speed in mm/sec according to the size standard in Microsoft Excel.

### Analysis of head-tail angles in freely behaving larvae

A custom written python script was used to analyze the tail bending angle of single freely swimming zebrafish larvae (https://github.com/MariusSeidenthal/zebrafish_angle_analysis). In short, this script applies a background correction to remove artifacts and detect the shape of the animal [51]. This is then skeletonized to a one-pixel line along the middle part of the body. The head is differentiated from the tail by analysis of surrounding pixel density in the binary image [52]. A single angle is then calculated between three points of the skeleton approximately corresponding to the tail tip, swim bladder and one point halfway between those.

### Electrophysiology

Zebrafish larvae 4 dpf were used for electrophysiological recordings. Experiments were performed in a darkened room with the light of the stereomicroscope (used for dissection) and on the microscope used for patch clamp recordings being covered with red filter foil to avoid prestimulation of bPAC as much as possible. A single zebrafish larva was transferred to bath solution (see below) containing tricaine (0.02%, MS222, Sigma Aldrich) for 1 minute to anesthetize it for dissection. Under a stereomicroscope (Stemi 2000, Zeiss, Germany) the larva was decapitated with a scalpel and then transferred to a recording chamber coated with Sylgard 184 (Dow Silicones Corporation, USA). It was fixed on its side with two tungsten pins through the notochord. The skin on the top was peeled off with a fine forceps to allow access to the underlying muscle cells. The recording chamber was then placed on an Axioskop 2 FS Plus microscope (Zeiss, Germany) equipped with DIC optics and a 40x water immersion objective. Electrodes for whole-cell muscle recordings were pulled to an outer diameter of 2–3 μm (resistance ∼5 MΩ) and lightly fire-polished using a Microforge MF-830 (Narishige, Japan). Bath solution (in mM): 134 NaCl, 2.9 KCl, 2.1 CaCl_2_, 1.2 MgCl_2_, 10 glucose, 10 Na-HEPES (pH 7.8). Intracellular solution (in mM): 120 KCl, 10 K-HEPES, 5 BAPTA (pH 7.4). Patch-clamp recordings were performed at room temperature using an EPC-10 amplifier (HEKA Elektronik, Germany). Data were acquired using the Patchmaster software (HEKA Elektronik, Germany) at a holding potential of -60 mV and filtered at 2,9 kHz. bPAC photostimulation was performed using a KSL-70 LED light (Rapp Optoelectronics, 470 nm), triggered by the Patchmaster software. Miniature EPCs (mEPCs) were analysed using Easy Electrophysiology software (https://www.easyelectrophysiology.com/).

### Statistical analysis

Data are shown as mean ± s.e.m. and n indicates the number of animals tested. Statistical significance between groups was determined by student’s t-test, Kruskal-Wallis test with Dunn post-hoc-test, 1-Way-ANOVA with Tukey multiple comparison of means or 2-Way-ANOVA with Tukey multiple comparison of means. The respective statistical test used is indicated in the figure legends. Generally mean ± SEM is shown unless otherwise stated. Data were analyzed and plotted with GraphPad Prism (GraphPad Software, version 8.02), Origin Pro 2023 (OriginLab) or the R statistical software environment (https://www.r-project.org/). Significance codes: ‘*’ p<0.05, ‘**’ p<0.01, ‘***’ p<0.001, ‘****’ p<0.0001).

## RESULTS

### Activation of bPAC in cholinergic motor neurons enhances locomotion activity

In order to analyze the effect of ChR2 and bPAC stimulation in embryonic and larval zebrafish motor neurons we generated transgenic lines expressing the respective optogenetic tools. The *mnx1* promotor [40, 53] has been described to be specifically active in primary and secondary motorneurons (PMNs and SMNs, respectively). We took two approaches: 1) we used the *mnx1* promotor fragment to directly express a fusion protein consisting of bPAC and EGFP (*Tg[mnx1En:bPAC-egfp]*); 2) we harnessed the *Tg[mnx1:Gal4]* driver line as a means to express ChR2 and bPAC from the *Tg[UAS:ChR2-P2A-dTomato]* and *Tg[UAS:bPAC-V2A-mCherry]* [35] transgenes, respectively. EGFP and dTomato fluorescence were clearly visible in PMN cell bodies and axons as early as 24 hours post fertilization (hpf) (**Fig. 1A**, left panels). Corresponding expression of the ChR2 and bPAC transgenes in PMNs and SMNs could be observed at 4 days post fertilization (dpf) (**Fig. 1A**, right panels).

**Figure 1:**
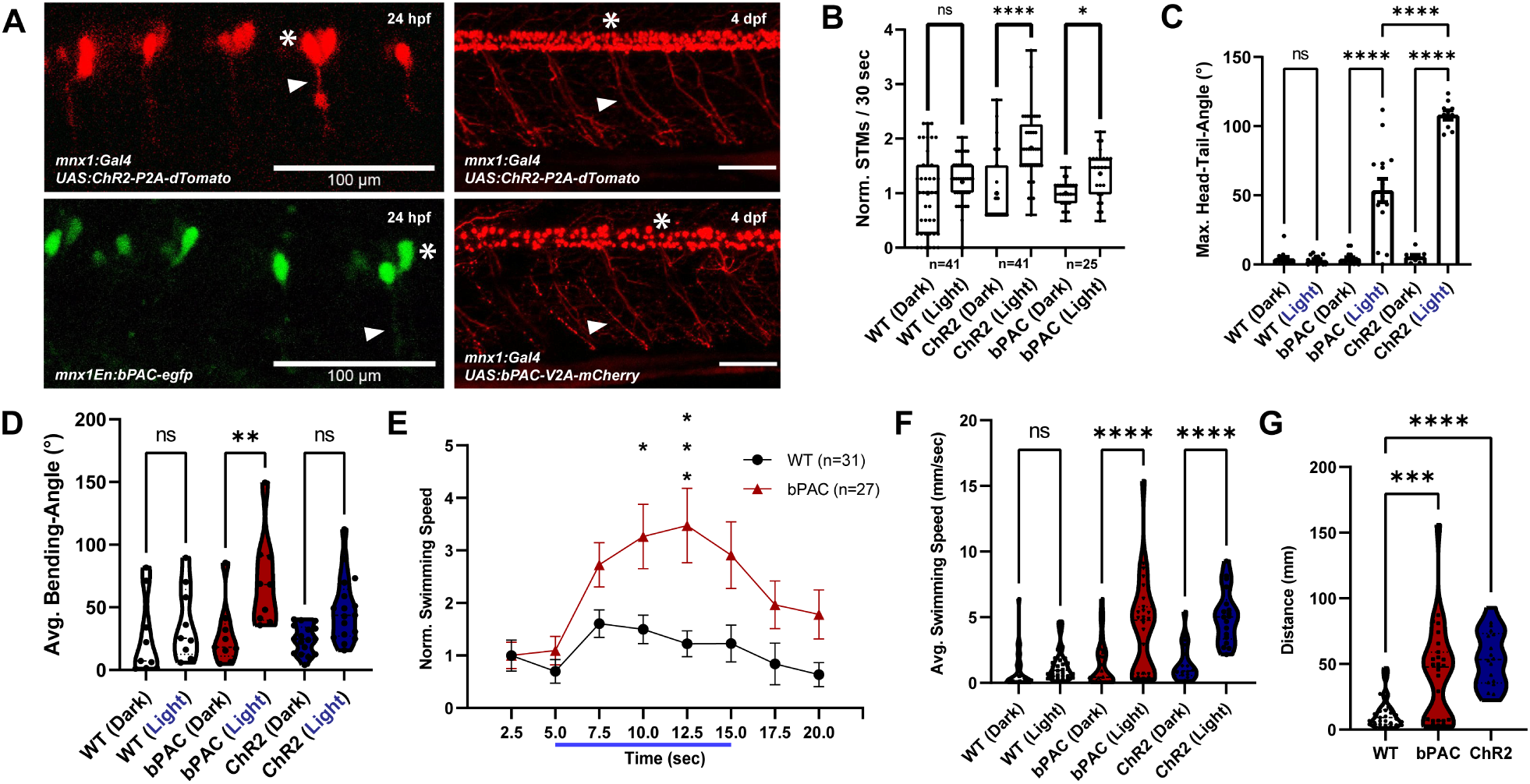
Expression and stimulation of ChR2 and bPAC in zebrafish motor neurons alters locomotion behavior. **(A)** Confocal images of Tg[*mnx1:Gal4*] / Tg[*UAS:ChR2-P2A-dTomato*] and Tg[*mnx1En:bPAC-egfp*] transgenic embryos (24 hpf) and of Tg[*mnx1:Gal4*] / Tg[*UAS:ChR2-P2A-dTomato*] and Tg[*mnx1:Gal4*] / Tg[*UAS:bPAC-V2A-mCherry*] double transgenic larvae (4 dpf). Asterisks mark motor neuron cell bodies, arrowheads motor axons. **(B)** Spontaneous (Dark) and evoked (Light) tail movements (STMs / ETMs) in 24 hpf embryos. **(C)** Head-tail-angles in immobilized larvae under dark and blue-light conditions. The maximum head-tail-angles were determined for wild type (WT), bPAC (Tg[*mnx1: Gal4*] / Tg[*UAS: bPAC-V2A-mCherry*]) and ChR2 (Tg[*mnx1:Gal4*] / Tg[*UAS:ChR2-P2A-dTomato*] transgenic larvae (4 dpf). **(D)** Bending angles of freely swimming larvae under dark and photostimulated conditions. The bending angles were determined for animals as in C. **(E)** Swimming speed was measured for individuals 4 dpf before, during and after blue light stimulation. **(F)** Average swimming speed with and without blue light stimulation. **(G)** Distance traveled during the photostimulation phase. In B, Kruskal-Wallis test with Dunn post-hoc-test., in C-F, Two-Way-ANOVA in G, One-Way-ANOVA, each with Tukey multiple comparisons of means. Statistical significance given as ****p<0.0001, ***p<0.001, **p<0.01, *p<0.05.

To test whether cholinergic motor neurons can be activated and neuromuscular signaling increased by elevated cAMP levels, behavioral assays of locomotion phenotypes were used. bPAC enzymatically generates cAMP upon activation by blue light of approximately 441 nm [33]. cAMP signaling, through PKA and its presynaptic targets, is expected to evoke increased neuronal activity, also in zebrafish motor neurons. Similarly, ChR2 depolarizes neurons during stimulation with blue light [54] and is thus expected to evoke motor activity [28, 55, 56]. Blue light stimulus frequency and intensity were optimized for the application of ChR2 and bPAC in accordance with previous work by others [27]. Motor activity in zebrafish embryos starts around 17 hpf with a side-to-side movement of the tail, a characteristic behavior called spontaneous tail coiling [57]. Since this behavior is mediated by motor neurons and interneurons in the spinal cord, we reasoned that it could be a possible early measure for optogenetically evoked neuro-muscular activity as well. Wild-type and transgenic zebrafish embryos at 24 hpf showed a similar baseline coiling rate under dark conditions (**Figs. 1B; S1A**). Blue light stimulation of wild-type animals did not evoke significant changes in behavior, however, for both, ChR2 or bPAC, blue light induced a significantly increased coiling rate in transgenic embryos, as compared to the respective condition without stimulus (**Figs. 1B; S1A**).

To analyze if activation of bPAC and, consequently, the increase of cAMP levels or the stimulation of ChR2 in motor neurons also triggers any locomotion changes in older zebrafish larvae, we performed further assays. Observing induced locomotion in partially immobilized larvae is an established way to determine if a certain input triggers e.g. an escape response or augmented swimming attempts [28, 58]. To this end, we measured the maximum deviation of the tail tip from the body axis, namely the angle between head and tail, in head-mounted animals, in the dark, and following blue light stimulation. Whole field illumination and respective activation of bPAC and ChR2 in all motor neurons led to large amplitude tail bends, while wildtype larvae did not exhibit any change in behavior and no escape responses (**Fig. 1C**). This indicates that bPAC as well as ChR2 can specifically activate motor neurons. We did not observe any seizure-or paralysis-like behavior in this assay. This indicates that MN activity was not induced by massive depolarization, but rather, that their membrane potential was elevated, such that intrinsic circuit activity evoked behavior more readily, yet in a coordinated way, to trigger muscle contraction. Based on the deeper bending angles, contractions were more pronounced for ChR2 stimulation.

Strong tail beating as evoked by MN stimulation via bPAC or ChR2 in immobilized larvae is expected to lead to an increased swimming speed in freely behaving animals. We established a method to record swimming behavior and quantify swimming parameters like the body bending angle and speed of larval zebrafish over time. Individual larvae (4 dpf) were placed in an agarose arena and subjected to an illumination protocol as described in the methods section. Wild type larvae exhibit a mild escape response upon illumination with bright blue light, as described previously [59]. However, a significantly stronger response was observed in bPAC and ChR2 transgenic larvae (Tg[mnx1:Gal4] / Tg[UAS:bPAC-V2A-mCherry], Tg[mnx1:Gal4] / Tg[UAS:ChR2-P2A-dTomato]). This became evident as increases in a) bending angles (**Fig. 1D**), b) swimming speed during the blue light stimulation phase (**Figs. 1E, F,; S1B-D**), and c) the distance which larvae cover during light stimulation (**Fig. 1G**). Particularly bPAC expressing animals showed a significant increase in locomotion behavior as compared to non-transgenic wildtype animals. Yet, some bPAC transgenic larvae did not respond to the blue light stimulus (**Fig. 1F, G**). Since we did not observe this for ChR2 transgenics (**Fig. 1F, G**) we speculated that this might be due to dark activity of bPAC [33], and adaptation of cAMP pathways.

Our findings in behavioral assays demonstrate that ChR2-induced depolarization of cholinergic motor neurons causes rapid behavioral changes (e.g. tail beating and swimming) which originate from enhanced muscle activity. bPAC stimulation evokes similar effects. These findings are in line with the hypothesis that an increased cAMP level in cholinergic cells might lead to increased rates of SV fusion and exocytosis of the neurotransmitter ACh.

### Neuropeptide signaling genes are expressed in larval motor neurons

We found that optogenetic cAMP generation induced locomotion behavior. Similar observations were made in motor neurons of other organisms. These could be linked to neuromodulatory activity, specifically signaling via neuropeptides, which in *C. elegans* was found to regulate the filling state of cholinergic SVs [14]. Previous studies revealed that cholinergic neurons in zebrafish express neuropeptides [60] and larval motor neurons contain dense core vesicles (DCVs), as visualized by electron microscopy [19]. However, it is not firmly established which roles these neuropeptides have. Given the cAMP effects on locomotion, we wondered if some of these neuropeptides might be involved in modulation of signal transmission at NMJs in zebrafish. Therefore, we analyzed whether key components of the neuropeptide machinery are expressed in motor neurons. Similar to other vertebrates, zebrafish neuropeptides are derived from longer precursor proteins, which undergo enzymatic post-translational processing in DCVs [21]. Publicly available single cell RNA sequencing data [61] and RNA *in-situ* hybridization showed that carboxypeptidase E (*cpe*), which is involved in biosynthesis of most neuropeptides and peptide hormones [20, 62, 63], is expressed in spinal cord neurons and motor neurons of 24 hpf embryos and 4 dpf zebrafish larvae, respectively (**Figs. 2A; S2**). By mass-spectrometry analysis of whole brain lysates, 62 neuropeptides originating from 34 different peptide precursors were identified in zebrafish [64], and expression of several neuropeptides, likely in the central nervous system, was validated. Furthermore, expression profiling of zebrafish neurons at 4 dpf by single cell RNA sequencing [61] identified a particular cluster of mRNA profiles (cluster #143) containing prominent marker genes for motor neurons like *mnx1* and *isl1* (**Fig. S2**). The same cell cluster expresses the neuropeptide tachykinin 1 (*tac1*) at high level and with high significance in comparison to the entire cell atlas (**Fig. S2**). Likewise, 24 hpf embryos exhibited strong expression of *tac1* mRNA in the brain and in a region of the spinal cord anatomically known to contain motor neurons (**Fig. 2A**). These data suggest that certain neuropeptides are expressed and processed into their active form in, and are likely also released from, motor neurons during developmental stages relevant for our analysis. Hence, in the following experiments we focused on the candidate genes *cpe* and *tac1*.

**Figure 2:**
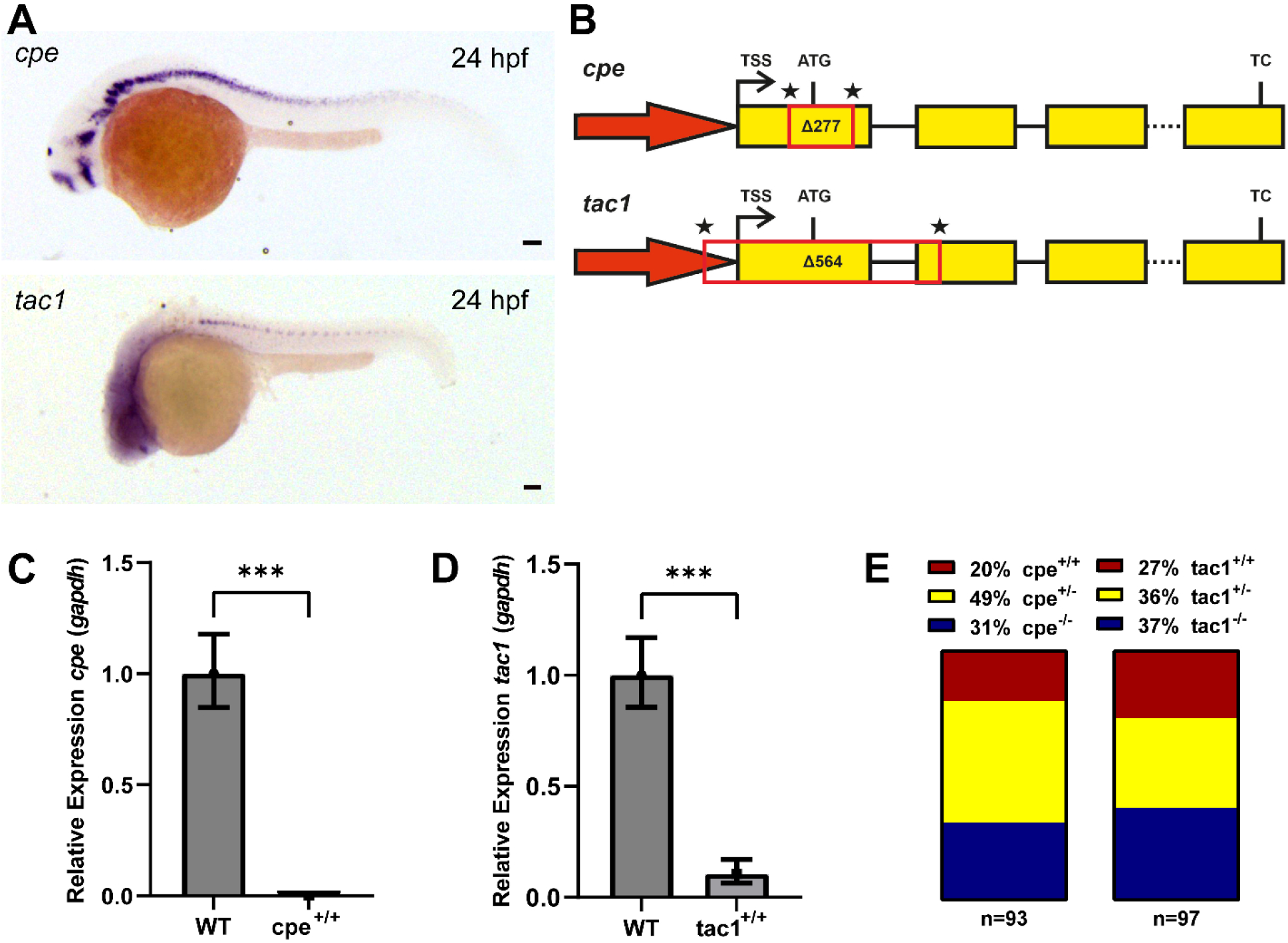
Key components of the neuropeptide signaling pathway are expressed in motor neurons. **(A)** Bright filed images of *in-situ* hybridization stainings for *cpe* and *tac1* mRNAs, respectively. Scale bars, 100 µm. **(B)** Schematic representation of *cpe* and *tac1* gene loci illustrating the CRISPR / Cas9 knockout strategy. Asterisks indicate sgRNA target sites. TSS: Transcriptional start site; ATG: Start codon; TC: Termination codon; PTC: Premature termination codon. **(C)** qPCR showing *cpe* mRNA levels relative to *gapdh*. **(D)** qPCR showing *tac1* mRNA levels relative to *gapdh*. **(E)** Percentage of wild type, heterozygous and homozygous genotypes within a population of 4 dpf *cpe* or *tac1* mutant larvae. In C, D, means ± standard deviation; Student’s t-Test. Statistical significance given as *p<0.05.

### Knockout of *cpe* and *tac1* by CRISPR-mediated gene deletion

To obtain knockouts, we used CRISPR / Cas9 technology, employing two sgRNAs for each target gene (**Fig. 2B**). For *cpe*, the resulting 277 bp deletion affects part of the 5’-UTR, the start codon and 202 bp of the coding sequence. For the *tac1* gene, we could induce a 564 bp deletion including part of the promotor, transcriptional start site, 5’UTR, start codon and the first 114 bp of the coding sequence together with the first splice site. In both the *cpe Δ277* and *tac1 Δ564* alleles, the respective mutant mRNA levels were significantly reduced as compared to wildtype animals (**Fig. 2C, D**) indicating a loss of the respective protein as well. *cpe*^-/-^ and *tac1*^-/-^ knockout larvae had no obvious morphological phenotype and developed normally until 4 dpf. Moreover, frequencies of genotypes observed from heterozygous breedings approximately matched the expected Mendelian inheritance patterns in 4 dpf larvae (**Fig. 2E**). *cpe^-/-^* knockout animals exhibit low survival rates during later development and die during juvenile stages, while *tac1^-/-^*knockout animals develop to adults without obvious phenotypes.

Hence, *cpe* and *tac1* homozygous knockout animals, as described above, were used for further experiments to analyze whether neuropeptides play a role in the cAMP-induced locomotion phenotype observed in photostimulated bPAC transgenic animals.

### Neuropeptides modulate signal transmission at zebrafish NMJs

Next, we subjected *cpe^-/-^* and *tac1^-/-^* animals to a swimming assay before and during bPAC stimulation, as described above. Basal locomotion speed during swimming before blue light stimulation was similar between wild type animals as well as bPAC transgenic animals of wild type and homozygous mutant siblings (**Fig. S3A, B**). When photostimulated, we found that homozygous *cpe* and *tac1* mutants swam significantly faster as compared to their wildtype siblings (more pronounced for *cpe* mutants; **Fig. 3A, B**). These findings could indicate that neuropeptides have a negative contribution to neuromuscular signaling, and thus, in their absence, higher locomotion speed increases are observed. Alternatively, neuropeptide signaling may positively influence synaptic transmission, and the observed increase of the behavioral response could be due to post-synaptic homeostatic scaling, compensating for a presynaptic deficit. *tac1* could encode one of the neuropeptides responsible for the observed phenotypes but is not solely responsible for the effects as observed in the *cpe* knockout animals.

**Figure 3:**
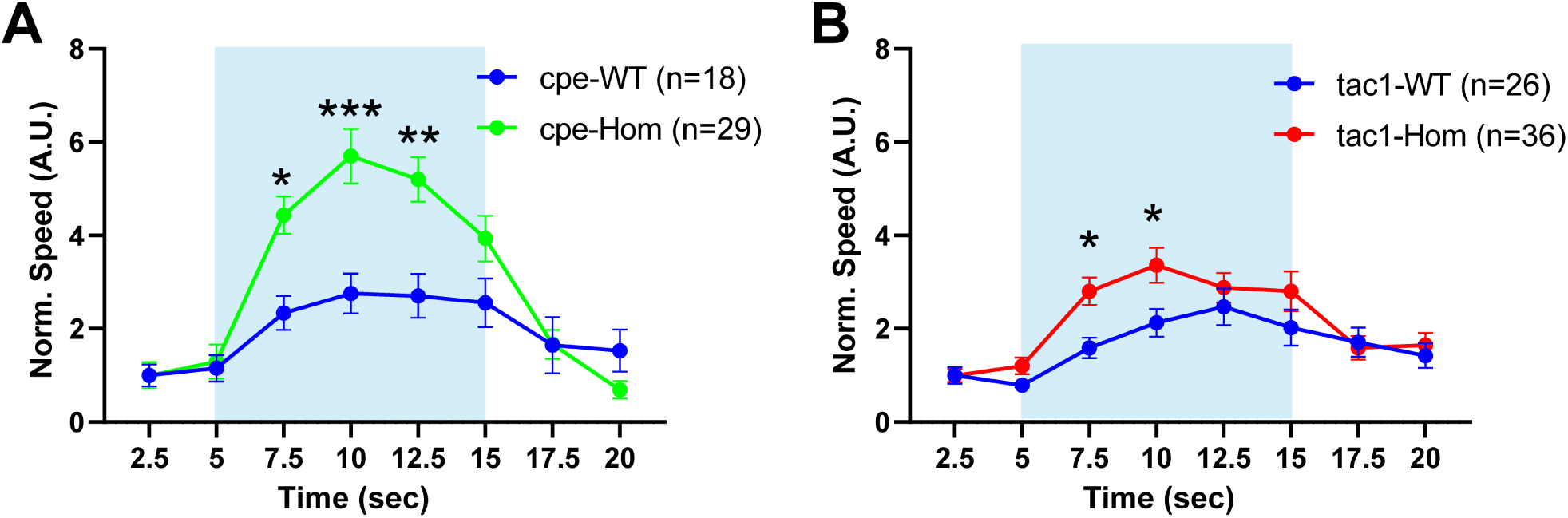
Enhanced locomotion behavior during bPAC stimulation in *cpe* and *tac1* knockouts. **(A)** *cpe* homozygous knockouts and wild type siblings (4 dpf) in the Tg[*mnx1:Gal4*] / Tg[*UAS:bPAC-V2A-mCherry*] double transgenic background were subjected to a swimming assay and swimming speed measured during blue light stimulation as indicated. Values are normalized to the first 2.5 seconds before illumination. Two-Way-ANOVA with Tukey multiple comparisons of means. **(B)** Swimming speed of *tac1* homozygous knockouts and respective wild type siblings (4 dpf) carrying the Tg[*mnx1:Gal4*] / Tg[*UAS:bPAC-V2A-mCherry*] transgenes before, during and after bPAC activation. Values are normalized to the first 2.5 seconds before illumination. Two-Way-ANOVA with Tukey multiple comparisons of means. Statistical significance given as ***p<0.001, **p<0.01, *p<0.05.

### Patch clamp recording from muscle cells shows altered transmission in *cpe^-/-^* mutants following bPAC photostimulation of MNs

To further characterize the mutants, and to explore the possible cause for their altered locomotion behavior, we turned to electrophysiology. Homozygous mutant animals were chosen for further analysis after genotyping on day 2 of development. We recorded from the superficial muscles of dissected larvae at 4 dpf, in voltage clamp mode, before, during and after 30 second blue light stimulation of cholinergic neurons expressing bPAC (**Fig. 4A**). During these experiments we found that a certain fraction of wild type animals did not show any effect during light stimulation in patch clamp recordings, even though expressing bPAC protein and responding to stimulation by a swimming response (judged by visual inspection; here we could not quantify this due to time constraints; note that we observed non-responders also in behavioral experiments; **Fig. 1F, G**). This might be due to damage of MNs or target muscle cells during the preparation. Since the number of animals which can be evaluated by electrophysiology is very limited we excluded the individuals described above from statistical analysis. We analyzed the rate of miniature excitatory post-synaptic currents (mEPSCs), which are evident as single current spikes of different amplitudes and are believed to reflect single SV fusion events. The basal rates of mEPSCs were smaller in wild type than in bPAC expressing animals (**Fig. 4B**), indicating an effect of bPAC expression, likely due to the known dark activity of bPAC, leading to elevated basal cAMP levels [33]. Also, *cpe^-/-^* animals expressing bPAC showed a lower basal mEPSC frequency compared to wild type and *tac1^-/-^* animals, though not reaching significance. When analyzed over the entire period of the experiment, normalized mEPSC frequency showed a significant increase during the blue light exposure in all bPAC-expressing strains, but not in wild type non-transgenic animals (**Fig. 4B, C**). This increase was significantly smaller in *cpe^-/-^* animals, compared to wild type, indicating a role of neuropeptides in regulating the rate of SV release or, possibly, mobilization. This neuropeptide, however, seems not to be encoded by *tac1*, as there was no difference observed in these mutants.

**Figure 4:**
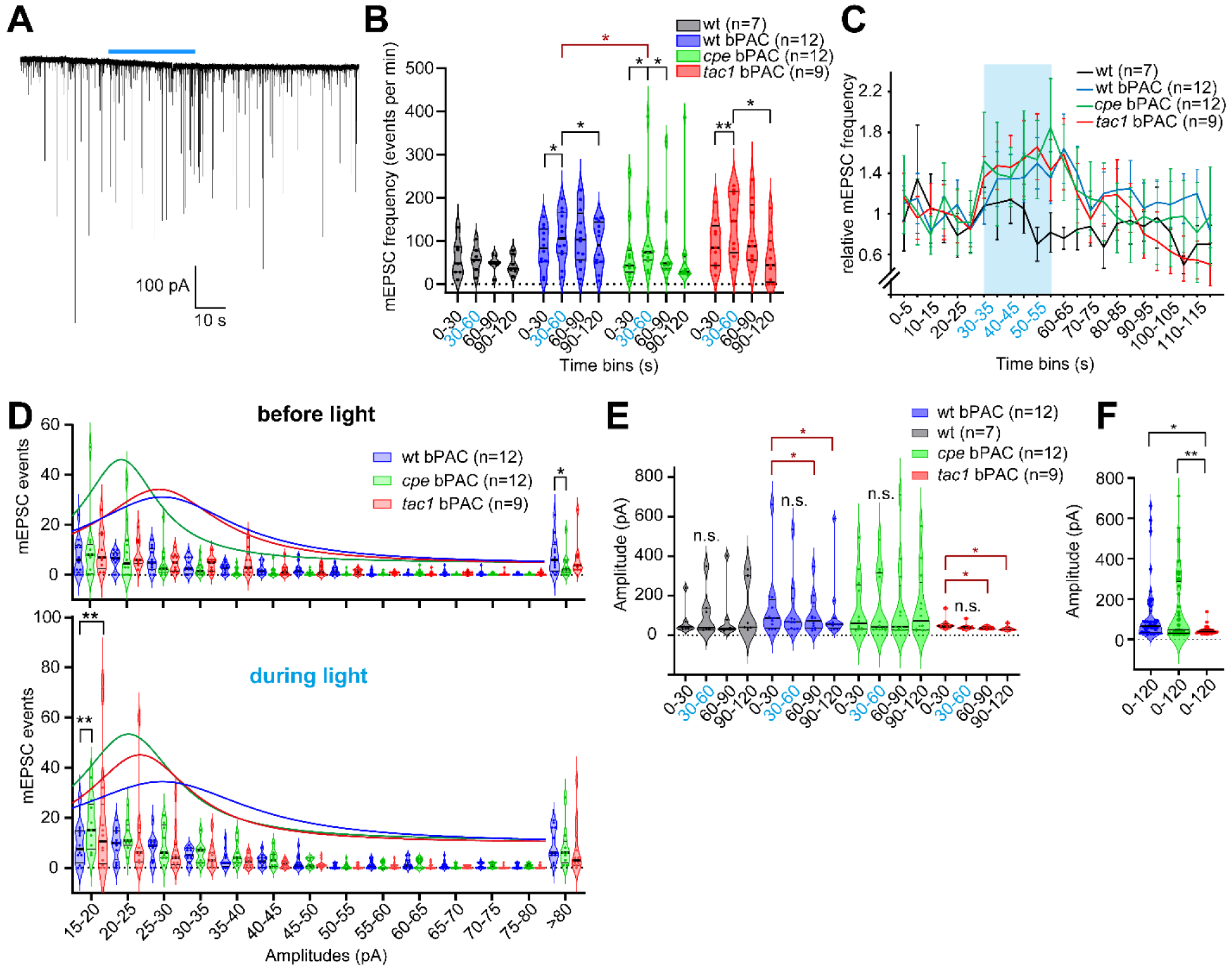
Post-synaptic recording from the NMJ following bPAC photostimulation demonstrates altered signaling in *cpe* and *tac1* knockout animals. A) Representative recording of mEPSCs before, during and after bPAC photostimulation (blue bar) in a wild type animal expressing bPAC in cholinergic motor neurons. **B)** mEPSC frequencies were analyzed in different time bins, as indicated, in wild type animals, as well as in bPAC-expressing animals, either wild type siblings, or *cpe^-/-^*or *tac1^-/-^* knockout animals. **C)** Normalized mEPSC rates analyzed in 5 s time bins, as indicated, in the respective animals as in B. Blue shade indicates time of blue light exposure. **D)** Number of mEPSC events of a given amplitude interval, as indicated, observed over a 30 sec interval before photostimulation (upper panel) or during the photostimulation (lower panel), for animals expressing bPAC, in wild type, or *cpe^-/-^*or *tac1^-/-^* knockout animals. **E)** mEPSC amplitudes were analyzed for each strain in 30 s intervals as indicated, before, during and after photostimulation. **F)** All mEPSC amplitudes across all intervals were analyzed for bPAC expressing animals (wildtype, or *cpe^-/-^* or *tac1^-/-^* knockouts, as indicated). In B, D-F, median and 25/75 quartiles (thick and thin black lines), min to max are shown. Two-way ANOVA, Tukey test. In B and E, red significance labels analyses using Fisher Test. Statistical significance given as **p<0.01, *p<0.05; n.s. – non significant.

Next, we analyzed the mEPSC amplitudes (**Fig. 4D-F**). Amplitudes exhibited a wide range between few dozen pA and up to 700 pA, in rare cases even >1 nA (**Fig. 4F**). This is difficult to reconcile, if single SV fusion events are causing them, but could for example be due to different release sites that are opposite of small or large clusters of postsynaptic nicotinic Acetylcholine Receptors (nAChRs), which mediate the currents underlying the mEPSCs. Moreover, different classes of motor neurons innervate the muscles, which may form neuromuscular junctions of different size. When we analyzed the mEPSC amplitudes before and during the photostimulation in the different genotypes, a similar distribution of frequencies and amplitudes became apparent (**Fig. 4D**). Before stimulation, *cpe^-/-^* and *tac^-/-^* animals showed a trend to smaller mEPSC amplitudes, which was significant during photostimulation, while wild type animals showed significantly more of the large mEPSC events (> 80 pA) compared to *cpe^-/-^* mutants. When analyzing mean amplitudes in the 30 s intervals before, during and after the light stimulus, there were significantly smaller amplitude events after the end of the light pulse in wild type and *tac1^-/-^* animals, but not in *cpe^-/-^* animals (**Fig. 4E**). Meanwhile, across all time periods, *tac1^-/-^* animals showed a significantly more uniform distribution of mEPSC amplitudes, ranging only from 20-100 pA (**Fig. 4F**).

In sum, the *cpe^-/-^* mutants, which are generally affected for neuropeptide signaling, showed fewer mEPSC events during photostimulation, more of the small and less of the large amplitude mEPSC events, while *tac1^-/-^* mutants had generally more uniform mEPSCs of moderate size. These findings may indicate that several neuropeptides with different modulatory effects could be involved in NMJ signaling. One of them may be inhibitory and lead to the reduced amplitudes after photostimulation, while the other could be promoting larger amplitude events; this latter peptide may be tac1, as these larger amplitudes were clearly absent in *tac1^-/-^* mutants.

### *cpe^-/-^* and *tac^-/-^* mutants express more postsynaptic nAChRs, possibly to compensate for altered neuronal ACh signaling

In the behavioral experiments, we observed an unexpectedly increased locomotion of *cpe* and *tac1* mutants in response to bPAC photostimulation, while these mutants were affected for presynaptic release of neurotransmitter. This could be explained by the absence of certain inhibitory neuropeptides in these mutant animals. Alternatively, there may be some homeostatic mechanism effective in the post-synaptic compartment that compensates for the reduced presynaptic cholinergic signal. One possible scenario of how this could be achieved involves reorganization or increased expression of nAChRs on the surface of muscle cells. To probe this hypothesis, we stained nAChRs using fluorescently labeled α-Bungarotoxin in 4 dpf larvae, i.e. in the same developmental stage the swimming assays were performed. α-Bungarotoxin binds specifically to the α-subunit of nAChRs, thus the resulting fluorescence signal reflects the amount and distribution of receptors on the muscle cell surface. Commonly, intense staining in large, connected clusters at the borders of the myotome are observed, oriented in dorso-ventral direction (**Fig. 5A**), as well as on the surface of the muscle cells in small discrete spots, likely representing sites of innervation by individual motor axons forming endplates [65, 66]. We found that in *cpe* and *tac1* mutants the mean number of small (rectangle in **Fig. 5B**) and large clusters (dashed rectangle in **Fig. 5B**) were not significantly different compared to wildtype animals (**Fig. S4A, B**). The size of the small clusters was significantly larger in *cpe^-/-^* mutants, while the size of the large clusters was not affected by loss of *cpe* or *tac1* (**Fig. S4C, D**). However, the fluorescence intensity of the small clusters was significantly increased in the *cpe* and *tac1* mutants (**Fig. 5C**) and strongly increased in large clusters for the *cpe* knockout (**Fig. 5C**). This might indicate that diminished cholinergic output from motor neurons, due to a loss of neuropeptides, is post-synaptically compensated by localizing more nAChRs to the muscle surface.

**Figure 5:**
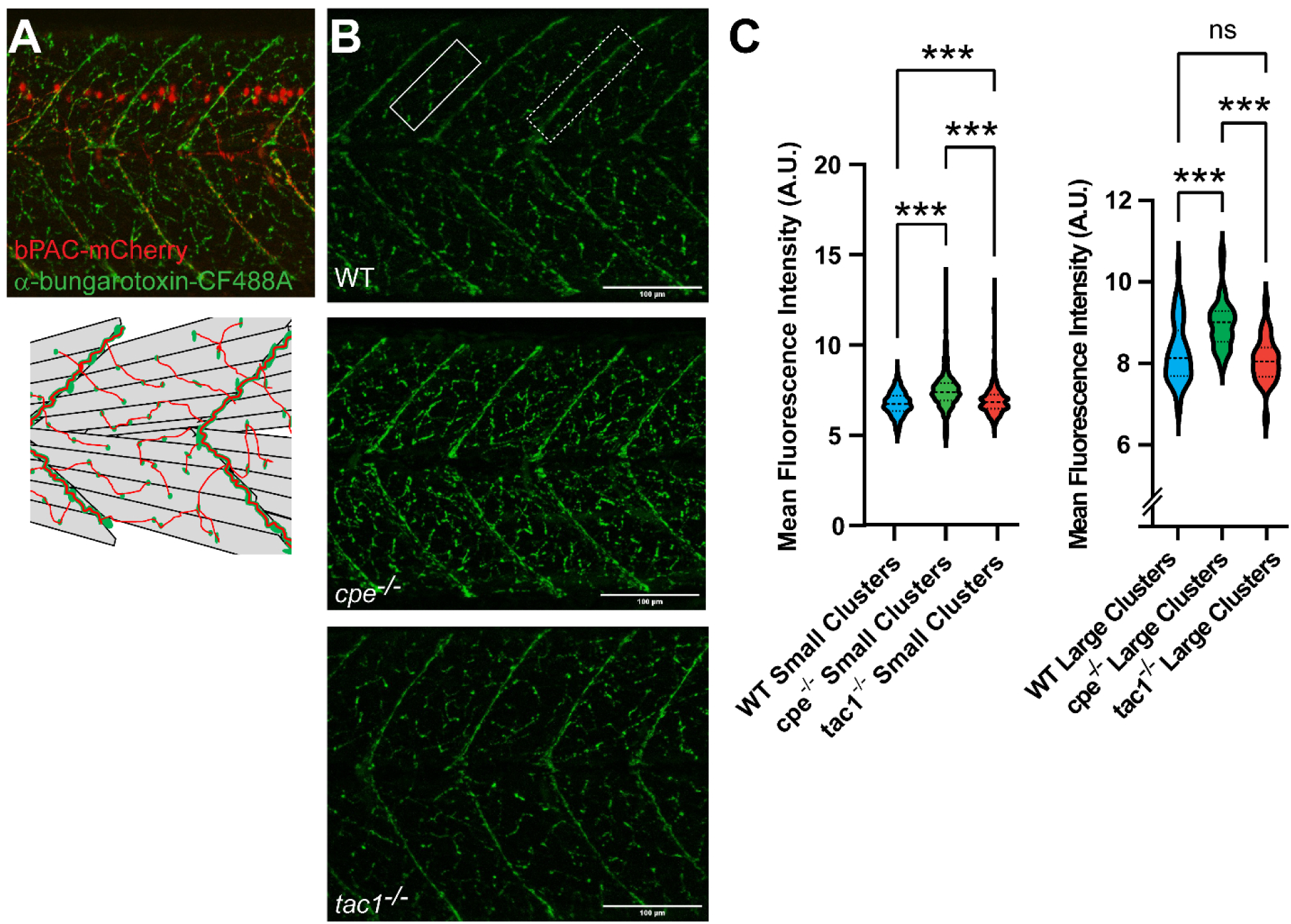
Abundance of nAChRs on muscle cells is altered in neuropeptide mutants. **(A)** Representative α-bungarotoxin staining at 4dpf (upper panel). nAChR clusters in green, bPAC expressing motor neurons in red. Diagram of skeletal muscle cells and clusters of nAChRs on the cell surface (lower panel). Large receptor clusters are assembled on somite boundaries, small receptor clusters are distributed in between. **(B)** α-bungarotoxin staining at 4dpf on wildtype, *cpe^-/-^* and *tac1^-/-^* animals. Rectangle in the upper panel represents an area containing small receptor clusters, dashed rectangle marks larger clusters at the end of muscle cells. **(C)** Quantification of fluorescence intensity of small and large receptor clusters in the respective zebrafish strains. Scale bars in B: 100 µm. One-Way-ANOVA with Tukey multiple comparisons of means. Statistical significance given as ***p<0.001; n.s. – non significant.

## DISCUSSION

The function of the neuromuscular junction is to excite muscle ensembles in a coordinated manner to enable movement of body parts, or for general locomotion. Different types of locomotion, i.e. slow, moderate swimming, or corrections of ongoing drifting movement, vs. rapid and vigorous swimming as part of an escape response, require different amounts of neurotransmitter release. Thus, motor neurons need to be able to secrete ACh in different quantities, which can be affected by different regulatory mechanisms: 1) Increasing depolarization causes increased action potential frequency, thus leading to fusion and mobilization of more SVs; 2) The level of SV mobilization and priming can affect overall transmitter release, as more SVs will fuse upon arrival of an action potential; 3) As observed in some species, the amount of released transmitter can also be mediated by the amount of ACh loaded into individual SVs. In the latter case, more release of ACh can be achieved from the same number of SVs, and this can be regulated in a very short time, using already existing SVs, and not requiring *de novo* SV formation at the synaptic endosome. In *C. elegans*, such a mechanism, i.e. acute loading of SVs with ACh, was observed in response to a rise in synaptic cAMP levels, using optogenetic stimulation [14]. Moreover, it was found that this regulation requires the release of neuropeptides from cholinergic motor neurons, thus likely acting in an autocrine fashion to influence the loading of SVs with ACh. In a natural setting, it is conceivable that exogenous signals, or intrinsic states of the animal associated with neuromodulatory signals in the motor nervous system (e.g. alertness and escape responses), may cause the increase in cAMP levels, to quickly upregulate neurotransmitter release in addition to the increase in firing rate. Also, more ACh release could be mediated even if the number of SVs becomes limiting, e.g. during periods of prolonged high activity. Our findings indicate that a similar mechanism may be present in zebrafish.

Neuropeptide signaling in zebrafish embryonal motor neurons was thus far mainly linked to developmental aspects of these cells. By electron microscopy, motor neuron terminals were shown to contain DCVs. Genes required for neuropeptide biogenesis, like carboxypeptidase E, as well as neuropeptide-encoding genes, have been identified in scRNAseq analyses of embryonal cells, clustering along with markers of motor neurons. By *in-situ* hybridization, we could show that two of these genes, *cpe* and *tac1*, are expressed in the spinal cord motor neurons. Knockouts could be obtained, eliminating formation of the respective mRNAs. We then analyzed behavioral and electrophysiological phenotypes, at basal level, and in response to optogenetic stimulation of the MNs using bPAC.

When motor neurons expressing bPAC were photostimulated, the animals showed a robust behavioral response, i.e. increased swimming speed, and deeper tail bending angles. In both mutants, *tac1*, and particularly, *cpe*, higher swimming speed was observed. This indicated not a malfunction, but rather a gain-of-function of the motor neurons. Alternatively, (postsynaptic) compensation in response to a presynaptic defect causing reduced release of ACh could affect larger increases in swimming speed following bPAC-induced neurotransmitter release.

Our electrophysiological recordings showed that bPAC activation induced an increased rate of mEPSCs, corresponding to single SV fusion events. The large variation in mEPSC amplitudes may be due to innervation of different size nAChR clusters, as we observed by bungarotoxin staining. Alternatively, large “compound vesicles” could have been formed from SVs first fusing to each other, and then being released in one large fusion event [67]. To address such a possibility, electron microscopy is required.

The mEPSC rate increase was not affected by the different mutations, apart from a somewhat lower release rate in the *cpe* mutant; this was not observed when the data were normalized. For the mEPSC amplitudes, we found that the usually broad distribution of amplitudes was much narrower in the *tac1* mutant, and the amplitudes were overall smaller. Wild type synapses produced more of the very large amplitude events, while *cpe* mutants showed more of the very small events, when photostimulated. Thus, there are some differences in ACh release, and/or ACh detection in the *cpe* and *tac1* mutants, that could underlie the observed behavioral phenotypes, either directly through the release of more (or less) transmitter, or by homeostatic compensatory alterations induced in these mutants. At present, we cannot exclude that a developmental defect caused the changes. However, morphologically we did not find obvious alterations in the appearance of NMJs, judged by the staining of postsynaptic nAChRs, though post-synaptic sites in *cpe* and in *tac1* mutants appeared to contain more nAChRs. Such upregulation may be a means to achieve the postsynaptic compensation.

One possible effect of neuropeptide signaling was indicated when we analyzed the mEPSCs in a time-dependent manner. Both wild type and *tac1* mutants showed smaller mEPSC amplitudes following bPAC photostimulation, which was not observed for *cpe* mutants. While we do not know if some adaptation or fatigue occurred that could explain these reduced amplitudes, the fact that they do not appear in a mutant presumably lacking all mature neuropeptides indicates that a neuropeptidergic regulatory mechanism may be involved. Since the motor neurons express several neuropeptides, there may be a balance of excitatory and inhibitory species, that is shifted towards the inhibitory side in the wild type, and not in place in the *cpe* mutant (i.e. no alteration in signaling occurs); in *tac1* mutants, the putative inhibitory signaling is still present, suggesting that *tac1* is not encoding this signal.

In sum, we showed that neuropeptidergic acute signaling occurs at the NMJ of larval zebrafish, or that neuropeptides shape NMJ signaling such that acute stimulation via cAMP causes different postsynaptic effects and exaggerated behavior. Whether cAMP triggers neuropeptide release acutely will have to be determined in the future, possibly using fluorescent neuropeptide release reporters to image release events *in-vivo* and real-time. However, such reporters have not yet been developed for zebrafish.

## Acknowledgements

We are grateful to Dr. Soojin Ryu (Johannes Gutenberg University, Mainz) for the ChR2-P2A-dTomato plasmid, and to Dr. Hernán Lopez-Schier and Dr. Amparo Acker-Palmer for zebrafish lines. We thank Franziska Baumbach, Hans-Werner Müller and Katharina Kuhlmeier for expert technical assistance and laboratory management. We acknowledge the animal care takers at the Goethe-University Biologicum, Ema Omeragić and Deeksha Gopinath Krishnamoorthy for help with animal husbandry and Dr. Bettina Kirchmaier for assistance with approval procedures. This work was supported by Deutsche Forschungsgemeinschaft (DFG) grant GO1011/19-1 and CRC1080-B02, to A.G., and by funds from Goethe University.

## Contributions

Conceptualization: H.D., A.G.

Methodology: H.D., J.F.L., A.G.

Software: H.D., M.S.

Validation: H.D., J.F.L., A.G.

Formal Analysis: H.D., A.G.

Investigation: H.D., J.F.L., M.B., A.G.

Resources: H.D., J.F.L., M.S., A.G.

Data curation: H.D., J.F.L., A.G.

Writing – Original Draft: H.D., J.F.L., A.G.

Writing – Review & Editing: H.D., J.F.L., A.G.

Visualization: H.D., A.G.

Supervision: A.G.

Project Administration: A.G.

Funding Acquisition: A.G.

---

**Figure S1:**
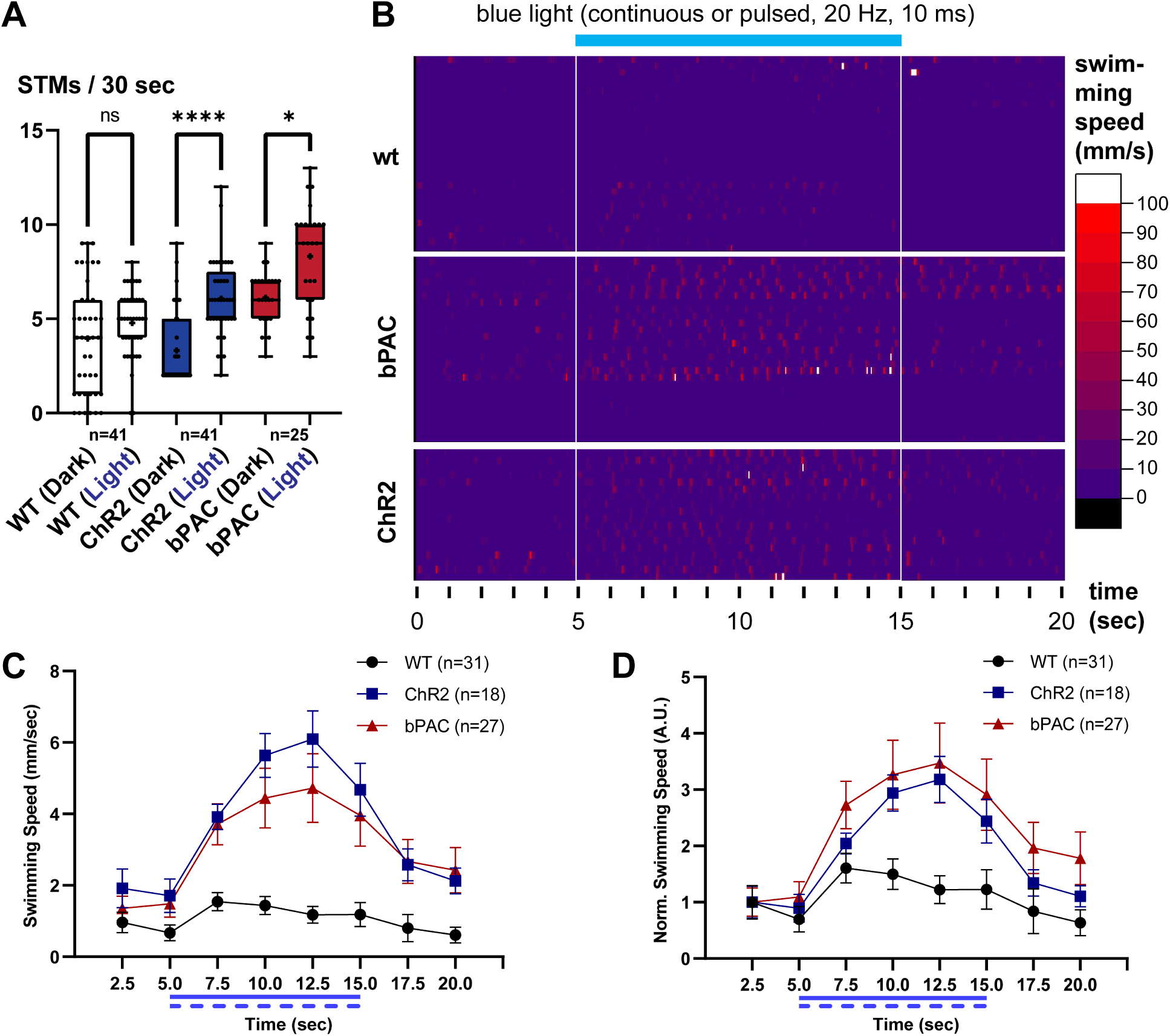
Locomotion during ChR2 or bPAC stimulation. **(A)** Spontaneous (Dark) and evoked (Light) tail movements (STMs / ETMs) in 24 hpf embryos. Kruskal-Wallis test with Dunn post-hoc-test. **(B)** Raster plot of swimming speeds, extracted from videos, at 30 Hz sampling frequency, color coded as indicated on the right, and for the respective animals expressing no transgene (wt) or bPAC or ChR2 in cholinergic motor neurons. Each line represents one animal. **(C)** Swimming speed as measured with and without light stimulation. Swimming speed was measured for wild type, ChR2 transgenic or bPAC transgenic individuals 4 dpf before, during and after blue light stimulation. **(D)** Swimming speed as measured with and without blue light stimulation and normalized to the first 2.5 second time interval. Animals as in C. Statistical significance given as ****p<0.0001, *p<0.05; n.s. – non significant.

**Figure S2:**
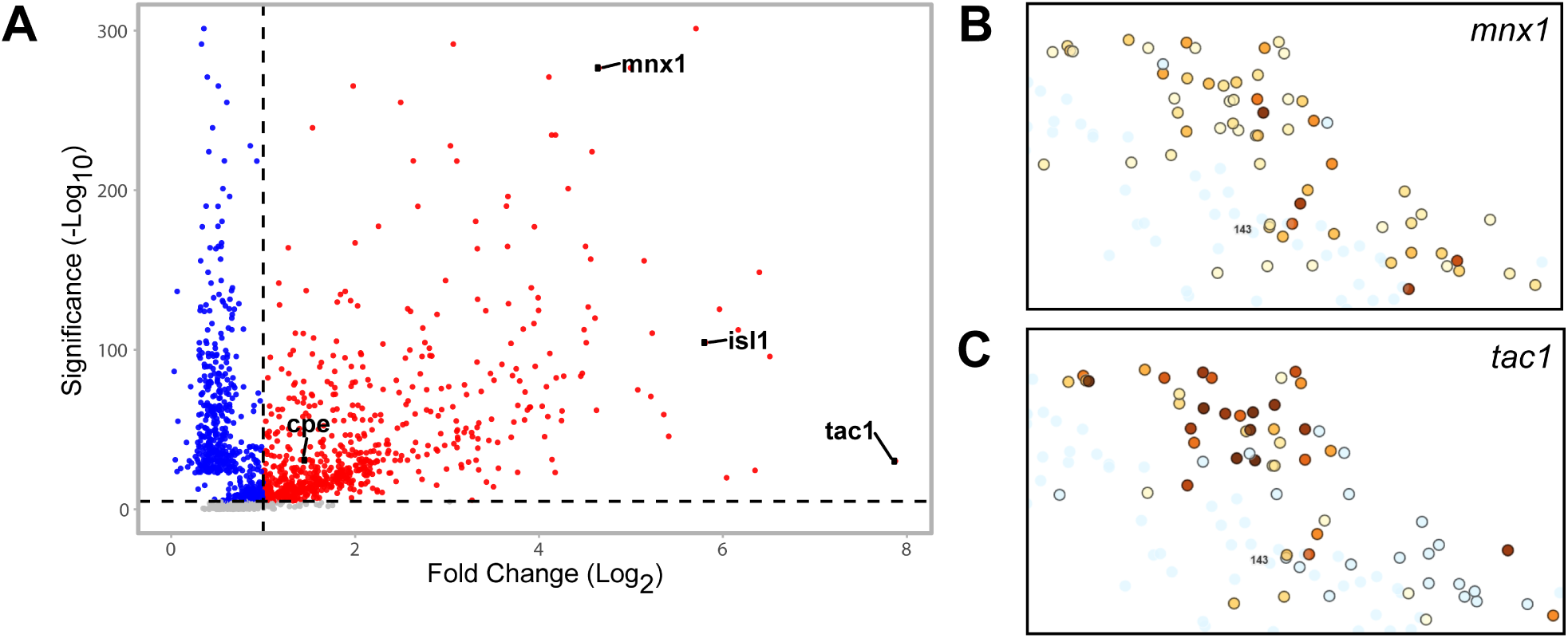
Genes encoding carboxypeptidase E *cpe* and the neuropeptide *tac1* are expressed in larval motor neurons. **(A)** Gene expression profile of the motor neuron cluster #143 obtained by single cell RNAseq. *mnx1* and *isl1*: Specific markers for motor neurons used to identify the cell type. **(B – C)** Magnified view of a motor neuron cluster shows high expression levels of the marker gene *mnx1* as well as the candidate gene *tac1*. Figures S2A – C adapted from [61] and the respective publicly available data sets (http://cells.ucsc.edu/?ds=zebrafish-dev).

**Figure S3:**
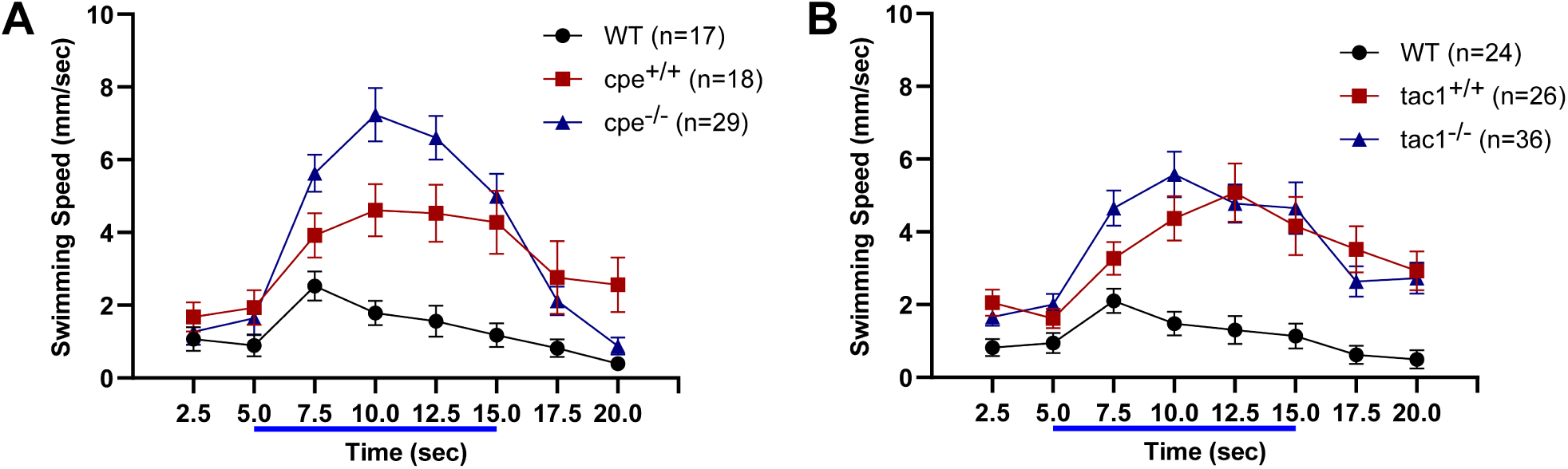
Locomotion behavior of *cpe* and *tac1* mutants. **(A)** Swimming speed was measured for wild type animals as well as *cpe* homozygous knockouts and wild type siblings (4 dpf) in the Tg[*mnx1:Gal4*] / Tg[*UAS:bPAC-V2A-mCherry*] transgenic background. **(B)** Swimming speed of wild type, *tac1* homozygous knockouts and respective wild type siblings (4 dpf) carrying the Tg[*mnx1:Gal4*] / Tg[*UAS:bPAC-V2A-mCherry*] transgenes before, during and after bPAC activation.

**Figure S4:**
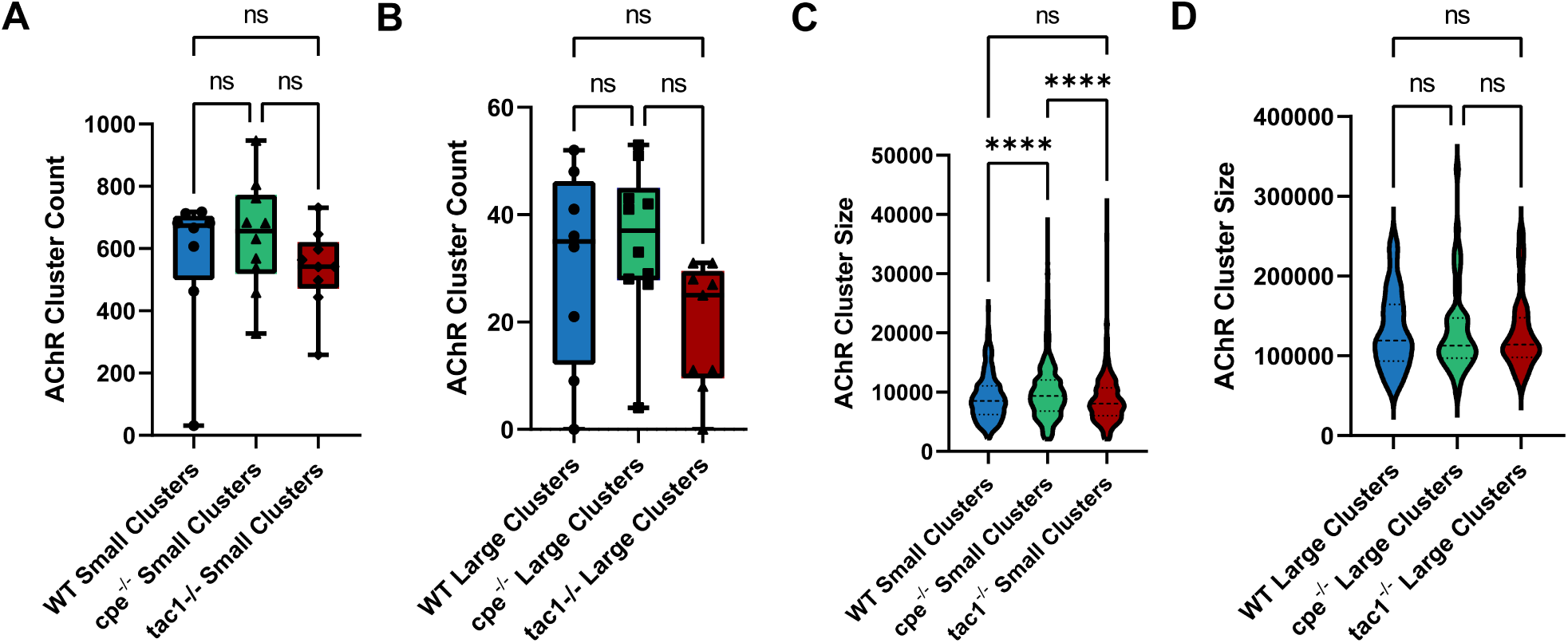
Analysis of nAChR cluster properties on muscle cells. **(A – B)** Quantification of the number of small and large receptor clusters in wild type animals and *cpe* or *tac1* homozygous mutants. **(C – D)** Quantification of small and large receptor cluster sizes in wild type animals and *cpe* or *tac1* homozygous mutants. One-Way-ANOVA with Tukey multiple comparisons of means. Statistical significance given as ****p<0.0001; n.s. – non significant

